# Syngeneic adipose-derived stromal cells modulate the immune response but have limited persistence within decellularized adipose tissue implants in C57BL/6 mice

**DOI:** 10.1101/2024.10.11.617847

**Authors:** John T. Walker, Tyler T. Cooper, Joy Dunmore-Buyze, Fiona E. Serack, Courtney Brooks, Aaron Grant, Maria Drangova, Gilles Lajoie, Gregory A. Dekaban, Lauren E. Flynn

## Abstract

The delivery of adipose-derived stromal cells (ASCs) on cell-instructive decellularized adipose tissue (DAT) scaffolds is a promising strategy for stimulating host-derived soft tissue regeneration. However, a better understanding of the mechanisms through which ASCs modulate regeneration *in vivo* is needed to harness these cells more effectively. In this study, DAT scaffolds, both with and without seeded syngeneic DsRED^+^ mouse ASCs, were implanted into immunocompetent C57BL/6 mice. Downstream analyses focused on assessing donor ASC persistence and phenotype, as well as the effects of ASC seeding on host macrophage polarization and the perfused host vascular network. Notably, most donor ASCs were cleared from the scaffolds by 2 weeks. Mass spectrometry-based proteomics indicated that the transplanted ASCs maintained their pre-implantation phenotype up to 1 week *in vivo*, suggesting that the cells were not undergoing programmed cell death. A higher fraction of the infiltrating host macrophages expressed CD68 and Arginase-1 in the ASC-seeded implants up to 1-week post-implantation. Interestingly, a small population of phagocytic macrophages, identified by uptake of DsRED protein, was present in the DAT implants in the first 2 weeks and showed enhanced expression of CD68, Arginase-1, and CD163, along with reduced expression of iNOS. MicroCT angiography revealed a similar perfused vessel network in the seeded and unseeded groups at 4- and 8-weeks post-implantation. Overall, seeding with syngeneic ASCs modulated the host macrophage response to the DAT bioscaffolds at early timepoints, but did not impact long-term regenerative outcomes, potentially due to the rapid clearance of the donor cell population in this model.

## Introduction

While adipose tissue shows a remarkable level of plasticity regulated by energy supply and demand^1,2^, it also has a limited capacity for self-renewal following tissue injury^3^. Current strategies to repair soft tissue defects include dermal fillers, which can temporarily augment volume but require repeated treatments, and autologous fat grafting, which can offer a more permanent solution, but is limited by variable longevity and volume retention^4,5^. Recognizing these limitations, there has been interest in the development of tissue-engineering approaches to promote more predictable and stable adipose tissue regeneration.

Cell-based strategies applying adipose-derived stromal cells (ASCs) in combination with biomaterials have shown promise for tissue engineering, with the ASCs contributing to regeneration through their paracrine activity^6^. Within native adipose tissue, ASCs play critical roles in the repair of neighboring tissues following injury^7^, as well as in self-renewal during homeostasis^8,9^. When applied as a therapeutic cell source, ASCs can modulate the immune response by secreting factors such as IL-6, M-CSF, and PGE2, which can promote macrophage polarization toward a more regenerative M2-like phenotype^10–14^. Similarly, ASCs can support implant integration by promoting localized angiogenesis through paracrine-mediated mechanisms. More specifically, *in vitro*, ASCs have been shown to express angiogenic growth factors including PDGF, FGF, VEGF and HGF^15,16^ and can promote endothelial cell tubule formation^17,18^. These findings have also translated *in vivo*, where ASC-seeded biomaterials promote increased CD31^+^ vessel density relative to unseeded controls^13,19^.

In developing therapies, biomaterials can be designed to replace the lost adipose tissue volume and retain the therapeutic cells at the implant site, as well as functioning as cell-instructive matrices to help guide the regenerative response^20^. In particular, decellularized adipose tissue (DAT) is a strong candidate for this application as it mimics the structure, composition, and biomechanical properties of the native extracellular matrix (ECM), and has been shown to promote the differentiation of endogenous progenitors toward an adipogenic lineage^21–25^. Importantly from a translational perspective, human adipose tissue is abundantly discarded as surgical waste making it an accessible starting material. Moreover, lyophilized decellularized materials can be stored stably at room temperature for on-demand application^26,27^.

In order to design tissue-engineering strategies that can be successfully applied for volume augmentation in humans, a better understanding of the mechanisms involved in the regenerative process is needed in pre-clinical animal models that have a functional immune system. For example, it is well established that ASCs do not persist indefinitely at the implant site^28–31^, but it is unknown whether their loss is due to targeting by host immune cells, or due to limitations of oxygen and nutrient supply in a hostile environment. From a tissue-engineering perspective, these differential outcomes require different solutions. Similarly, it is important to assess how the implant environment affects donor ASC function. The interaction between ASCs and macrophages has been explored *in vitro*^14,32,33^, but it is largely unknown how these cells will interact in the complex *in vivo* environment of an implant site. Lastly, although angiogenesis is often linked to desirable outcomes, excessive inflammation and fibrosis are also associated with potent angiogenic responses, where the resultant vasculature is unstable and poorly perfused^34^. Therefore, it is necessary to characterize vascular regeneration using methods beyond standard characterization of endothelial cell abundance. In particular, micro- and macro-vascular perfusion are clinically relevant parameters that can provide insight into overall tissue health^35^.

In this study, DAT biomaterials with or without seeded syngeneic ASCs, were implanted into the inguinal region of immunocompetent C57BL/6 mice and analyzed at up to 8 weeks post-implantation. Donor ASCs expressed the red fluorescent protein DsRED to enable longitudinal tracking via flow cytometry. The DsRED^+^ donor cells were isolated from implants collected at days 3 and 7 post-implantation for proteomics assessment via mass spectrometry to gain insight into their phenotype over time relative to pre-implantation controls cultured on DAT scaffolds or tissue culture polystyrene (TCPS). In addition, flow cytometry analyses were performed to characterize the effects of ASC seeding on the phenotype of host macrophages infiltrating the implants over time. These analyses were complemented by characterization of the perfused vascular network within and surrounding the implants at 4- and 8-weeks post-implantation using a micro-computed tomography-based angiography approach.

## 1. Methods

### 2.1 Murine ASC isolation

All animal studies were performed with approval from the Animal Care Committee at Western University (AUP #2019-079) and followed Canadian Council on Animal Care (CCAC) guidelines. To enable long-term cell tracking, ASCs were isolated from B6.Cg-Tg(CAG-DsRED*MST)1Nagy/J mice that ubiquitously express DsRED. The mice were euthanized at 9-11 weeks of age by CO_2_ asphyxiation, followed by cervical dislocation. The inguinal fat pads from 2-4 mice of the same sex were extracted and pooled, rinsed in ice-cold phosphate buffered saline (PBS), and further processed by mincing. The tissue was then digested with 4 mg/mL type IV collagenase (Gibco) in DMEM/F12 media (Wisent) supplemented with 100 IU penicillin and 100 µg/mL streptomycin (P/S; Wisent) for 75 minutes at 37°C, at 100 rpm on a rotating platform. Following digestion, one volume of DMEM/F12 with 10% FBS and P/S was added to inactivate the collagenase. The sample was then centrifuged at 500 x g for 5 minutes, resuspended by vortexing, and centrifuged again. The lipid layer was aspirated, along with the digestion solution. The cell pellet was rinsed twice in DMEM/F12 with 10% FBS and P/S, and seeded in a single T75 culture flask. Cells were allowed to adhere for 48 hours, and the flasks were then rinsed to remove non-adherent cells and debris. The media was changed every 48-72 hours, and the cells were passaged every 48-96 hours when they had reached ∼90% confluence. After the initial seeding (P0), subsequent passages were re-plated at 500,000-600,000 cells per T75 flask. Cells were isolated at the end of P2 to be used for scaffold seeding.

### 2.2 Adipose tissue decellularization and scaffold preparation

Surgically-resected human subcutaneous adipose tissue was obtained with informed consent (HSREB 105426) and decellularized using a detergent-free process, as previously described^21^. Briefly, the adipose tissue was processed through freeze-thaw cycles, enzymatic digestion, and polar solvent extraction to remove cellular and lipid content. Following decellularization, the DAT was lyophilized and stored until needed. Scaffolds were prepared by cutting the DAT into 6 ± 1 mg (dry weight) fragments. These scaffolds were then decontaminated and rehydrated with a gradient series of ethanol diluted in PBS starting with 100% and followed by 90%, 70%, 35% ethanol in PBS. Rehydrated scaffolds were rinsed 3 times in PBS to remove residual ethanol. Prior to seeding, the scaffolds were equilibrated in ASC culture media at 37°C, 5% CO_2_ for 24 hours.

### 2.3 Scaffold seeding

ASCs were harvested from culture plates using 0.25% Trypsin/ 0.1% EDTA and counted with a hemocytometer. Each scaffold was seeded with 1.0 x 10^6^ cells, in a 4 mL volume of DMEM/F12 with 10% FBS and 1% P/S, within a vented 15 mL conical tube (CELLTREAT). These conical tubes were then placed at a 15° angle on a rotating platform at 50 rpm, and incubated at 37°C, 5% CO_2_ for 72 hours. Following dynamic seeding, the scaffolds were rinsed in PBS to remove non-adherent cells. Seeded scaffolds were then cultured statically in Prime-XV media (Irvine Scientific) for an additional 8 days, with media changes every 48-72 hours. Unseeded scaffolds were prepared alongside their seeded counterparts.

### 2.4 Inguinal implant model

Male and female C57BL/6J mice at 9-11 weeks of age were used as implant recipients in this study as outlined for each experiment. As a pre-operative analgesic, mice were injected with 2 mg/kg meloxicam at 30 minutes prior to surgery. The mice were then anesthetized with isoflurane. To prepare the surgical site, hair was shaved in the inguinal region, and the area was cleaned with chlorhexidine and alcohol. As a local anesthetic, 2 mg/kg of bupivacaine was injected at the surgical site. One incision (3-5 mm in length) was made in the skin on each side of the body. Donor-recipient sex-matched implants and unseeded controls were implanted contralaterally, each in proximity to their respective inguinal fat pad. The incisions were closed with 6-0 monocryl sutures, and the mice were provided with a subcutaneous saline bolus to maintain hydration. The following day, the mice received 1 mg/kg meloxicam for analgesia. Mice were monitored until endpoints ranging from 3-56 days.

### 2.5 Immunofluorescence Staining

To assess implant integration in terms of host cell infiltration, mice were euthanized at endpoint and the implants were excised within their surrounding tissues for analysis via immunofluorescence (IF) staining. The mice were euthanized by CO_2_ asphyxiation, followed by cervical dislocation. The samples were fixed overnight in 10% neutral buffered formalin and then prepared for cryopreservation through dehydration by subsequent washes in 15% and 30% sucrose in PBS. The tissue was then embedded in OCT for cryosectioning and subsequent immunostaining. Cryosections were prepared for labelling by washing in PBS with 1% sodium dodecyl sulfate, followed by PBS rinses and blocking in 10% donkey serum. Primary and secondary antibodies used in this study are listed in Supplemental Tables 1 and 2, respectively. Images were taken at 10X magnification on a LSM 800 confocal microscope (Zeiss) and stitched together. A minimum of 5 mice were observed for each stain at each timepoint, with tissues derived from at least one mouse of each sex per timepoint.

### 2.6 Fluorescence-activated cell sorting

To isolate the syngeneic donor ASCs from the DAT implants collected at endpoint, a fluorescence-activated cell sorting approach was used. Additional control samples were also processed following culture on TCPS, prior to scaffold-seeding, as well as from scaffolds that were seeded, but not yet implanted into mice. To prepare single cell suspensions, the scaffolds were extracted, manually minced, and digested in 4 mg/mL type IV collagenase (Gibco) in DMEM/F12 with 1% P/S for 2 hours at 37°C on a rocking platform set to 100 rpm. One volume of DMEM/F12 with 10% FBS, 1% P/S was then added to the digested scaffolds. The samples were centrifuged at 500 x g, resuspended by vortexing, and centrifuged again. The cells were then resuspended and filtered sequentially through 100 µm and 40 µm cell strainers, to obtain a single-cell suspension. The isolated cells were first labeled with SYTOX ™ Blue (Thermo Fisher Scientific) using a concentration of 1 µM to identify dead cells. Next, to separate the donor ASCs from host macrophages, the macrophages were labelled with a CD11b antibody. Samples were sorted on a BD FACSAria III sorting cytometer. The antibody, fluorophores, lasers, and filter sets used in this experiment are provided in Supplemental Table 3.

Live cells were first gated based on the absence of SYTOX ™ Blue. Singlets of appropriate size were then selected based on forward- and side-scatter parameters. ASCs were selected based on expression of DsRED and absence of CD11b labelling. Following collection, the sorted samples were rinsed 3 times in PBS, pelleted and stored at −80°C until needed for mass spectrometry analysis.

### 2.7 Mass spectrometry

To assess how the ASC population changed following implantation, proteomics assessment was performed via biological mass spectrometry. Due to the declining numbers of this population over time, this analysis was performed only up to 7 days post-implantation, when the ASCs were most abundant. Controls for this experiment included ASCs from the same donors cultured on TCPS prior to scaffold seeding, as well as those that had been seeded onto scaffolds, isolated prior to implantation. All groups underwent isolation via FACS as outlined above. Female donor cells implanted into female recipient mice were exclusively used for this experiment.

#### 2.7.1 Ultra-Performance Liquid Chromatography Tandem Mass Spectrometry

ASCs were lysed in an 8 M Urea, 50 mM ammonium bicarbonate (ABC), 10 mM dithiothreitol (DTT), 2% SDS lysis buffer using tip-probe sonication (10 X 0.5s pulses; Level 1) (Fisher Scientific, Waltham, MA). Protein was quantified by 660 nm spectrometry using detergent-compatible reagents. Approximately 10-25 µg of protein was reduced in 10 mM DTT for 30 minutes at room temperature, alkylated in 100 mM iodoacetamide for 30 minutes at RT in the dark, and precipitated in chloroform/methanol/water (1:4:3). On-pellet in-solution protein digestion was performed in 100 µL 50 mM ABC (pH 8) by adding Trypsin/LysC (Promega, 1:50 ratio) to precipitated ASC proteins. ASC proteins were incubated at 37°C overnight (∼18h) in a ThermoMixer C (Eppendorf) at 900 rpm. An additional volume of LysC (Promega, 1:100 ratio) was added for ∼3 hours before acidifying to pH 3-4 with 10% formic acid. Approximately 1 µg of peptides (estimated by bicinchoninic acid assay) were initially loaded onto an ACQUITY UPLC M-Class Symmetry C18 Trap Column, 5 µm, 180 µm x 20 mm and trapped for 4 minutes at a flow rate of 10 µL/min at 99% A/1% B. Peptides were separated on an ACQUITY UPLC M-Class Peptide BEH C18 Column (130Å, 1.7µm, 75µm X 250mm) operating at a flow rate of 300 nL/min at 35°C using a non-linear gradient consisting of 1-7% B over 1 minute, 7-23% B over 180 minutes and 23-35% B over 60 minutes before increasing to 98% B and washing. Settings for data acquisition on Q Exactive Plus are outlined in Supplemental Table 4.

#### 2.7.2 Proteomic Data Analysis

MS raw files were searched in MaxQuant (1.5.8.3) using the Mouse Uniprot database (updated November 2020 with 40,550 entries). Missed cleavages were set to 3 and I=L. Cysteine carbamidomethylation was set as a fixed modification. Oxidation (M), N-terminal acetylation (protein), and deamidation (NQ) were set as variable modifications (max. number of modifications per peptide = 5) and all other settings were left as default. Precursor mass deviation was left at 20 ppm and 4.5 ppm for first and main search, respectively. Fragment mass deviation was left at 20 ppm. Protein and peptide FDR was set to 0.01 (1%) and the decoy database was set to revert. The match-between-runs feature was utilized across all sample types to maximize proteome coverage and quantitation. Datasets were loaded into Perseus (2.0.6.0) and potential contaminants were removed. Of the remaining 4621 total identified proteins, those which were detected in at least 70% of the samples in at least one group (TCPS, DAT, day 3 implants, day 7 implants) were analyzed further. Samples with a limited proteomic depth were removed from further analysis using a limit of 2000 unique proteins identified (1 of 25 samples removed at this step). To be able to perform imputation independently for each group, the total list of proteins was further refined to include only proteins that were detected in at least one sample of each group. The remaining dataset included 2585 unique proteins and 24 total samples split up into the 4 groups (8 TCPS samples; 7 DAT samples; 5 day 3 implant samples; 4 day 7 implant samples). Imputation was performed using the missing forest approach in R with the missForest package (version 1.5). MaxQ LFQ intensity data were then transformed into relative protein abundance values with a 2^x^ transformation. These data were further processed in R to identify proteins that were significantly altered between groups. A linear mixed effect model was run for each protein using the nlme package (version 3.1-165) where the grouping variable was used as the fixed effect, and the ASC donor used as the random effect. P-values were tabulated and adjusted for multiple comparisons using a false discovery rate correction. Proteins with a false discovery rate adjusted p-value <0.05 and a log2 fold change >|1| were considered to be significantly differentially expressed. Hierarchical clustering and principal component analysis were performed on Z-value normalized protein abundance values.

### 2.8 Flow cytometry

To assess donor ASC persistence and host macrophage phenotype, the implants were excised at endpoint and processed for analytical flow cytometry. The chosen panel includes the following markers: CD11b, a marker for monocytes and macrophages; F4/80, a marker for macrophages; CD68, a widely expressed macrophage marker associated with phagocytic and lysosomal activity; CD163, a scavenger receptor expressed by M2 polarized macrophages; Arginase I, a marker for M2 polarized macrophages; and iNOS, a marker for M1 polarized macrophages.

The implants were processed as described above to prepare single cell suspensions. Cells were first labelled with LIVE/DEAD™ fixable Aqua dead cell stain (Thermo Fisher Scientific). Blocking was performed using CD16/CD32 blocking solution (BD) as per the manufacturer’s instructions. Antibodies for extracellular macrophage targets F4/80, CD11b, and CD163, were added for 30 minutes on ice. The cells were then fixed in 4% paraformaldehyde for 10 minutes at room temperature, followed by permeabilization using the FoxP3 Fixation/Permeabilization kit (eBioscience), as per the manufacturer’s instructions. Permeabilized cells were then labelled with antibodies for the intracellular macrophage targets CD68, Arginase I, and iNOS for 30 minutes on ice. The samples were rinsed, incubated in 4% paraformaldehyde for 5 minutes, followed by further rinsing in PBS with 5% FBS, and left at 4°C overnight, to be run within 24 hours on a BD LSRII flow cytometer. Dilutions, lasers, and filter sets used in this experiment are provided in Supplemental Table 5. Single color controls were made with CompBead Plus anti-rat compensation beads (BD) for antibodies, or from cells for the dead cell stain and DsRED controls. Fluorescence-minus-one (FMO) controls were used to set gates.

Samples were first gated for the live cell population based on LIVE/DEAD™ labelling. Single cells of appropriate size were then selected using forward- and side-scatter parameters. ASCs were identified as DsRED^+^, F4/80^−^, CD11b^−^ cells. Macrophages were selected based on expression of both F4/80 and CD11b, and could be further broken down into DsRED^+^, and DsRED^−^ populations based on their uptake of the DsRED protein derived from implanted ASCs. Of the 30 total mice used for this experiment, 3 mice were excluded from analysis due to abnormally large cell yields from extraction, indicative of contaminating cell populations, potentially from blood while harvesting. After exclusion, 5-6 mice were analyzed at each timepoint with a minimum of 2 of each sex.

### 2.10 Angiography

At endpoint, mice were injected with 1 U/g of heparin intraperitoneally to prevent clotting and then injected with 150 mg/kg ketamine with 7.5 mg/kg xylazine for anesthesia. The chest cavity was surgically opened and the right atrium cut to euthanize the mouse by exsanguination.

Immediately following euthanasia, heparinized saline (2-3 mL of 1 U/mL) was injected into the left ventricle to flush the vasculature. The descending aorta was then canulated and heparinized saline was continuously flushed through the vasculature at 100 mmHg for 5 minutes. The mice were then perfused through the canulated aorta with a silicone elastomer-based contrast agent containing erbium oxide nanoparticles (ErNP) based on the formulation by Tse *et al.* (2017)^36^. Mice were left to perfuse at 100 mmHg using a gravity-fed perfusion system for at least 2 hours to enable the elastomer to polymerize and were then preserved in 10% neutral buffered formalin until scanned. High resolution scans were acquired with a GE eXplore Locus scanner (GE Healthcare, London, ON). Projection images were acquired at 80 kVp, 450 µm with an exposure of 4500 ms. In total, 900 projections were acquired at 0.5° increments. Images were then reconstructed with a 20 µm voxel size and voxel scalar values were normalized to Hounsfield units.

The scans were analyzed in 3D slicer software (http://www.slicer.org). Vessels were segmented by thresholding the scalar value to 140 units and noise was removed with the “smoothing” function at 3 x 3 x 3 voxel dimensions. Scaffolds were manually segmented using the “paint” and “fill between slices” functions followed by the “smoothing” function. In total, 5-6 mice were scanned per timepoint with a minimum of 2 of each sex per timepoint.

## 2. Results

### 3.1 ASCs are mostly cleared from the implant site within two weeks of implantation

Preparation of ASC-seeded scaffolds through a dynamic seeding process followed by a static expansion phase in Prime-XV serum-free media resulted in broad distribution of the ASCs throughout the scaffolds, with a qualitatively enhanced cell density in the more peripheral regions of the scaffolds (Figure 1a). To assess the *in vivo* persistence of the donor ASCs, scaffolds were isolated at 3-, 7-, 14-, 28-, and 56-days post-implantation and digested for analysis via flow cytometry (Figure 1b). As a fraction of the total cell isolate, the donor DsRED^+^ ASCs decreased from 50.3% (±11.8%; n = 6) at day 3 to 3.74% (±0.86%; n = 5) at day 14 post-implantation, suggesting relatively rapid clearance of the population (Figure 1c).

**Figure 1:**
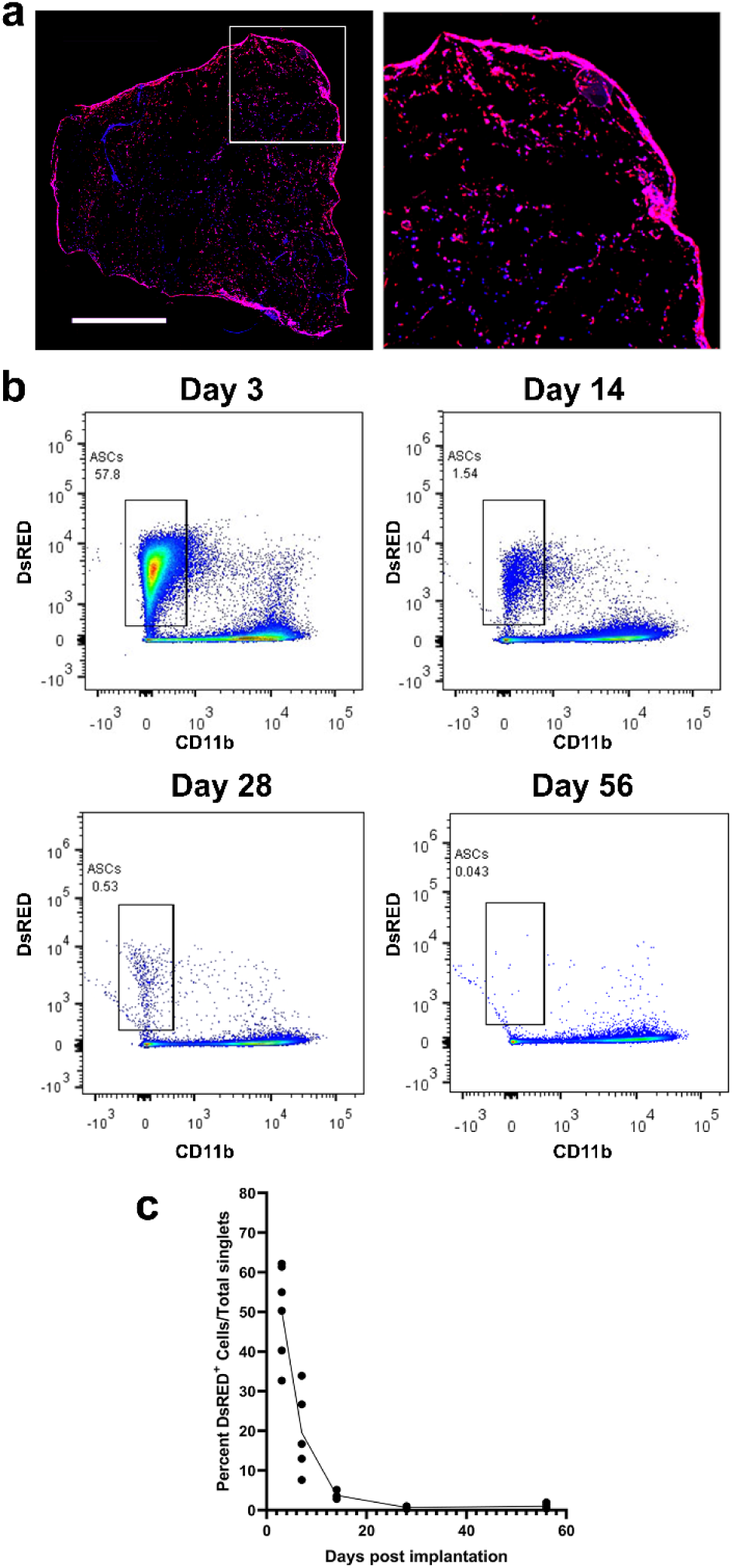
Syngeneic DsRED^+^ ASCs implanted in DAT scaffolds into the inguinal region of immunocompetent C57BL/6 mice were mostly cleared from the scaffolds within two weeks of implantation. (a) Prior to implantation, DsRED^+^ ASCs (magenta) were distributed throughout the DAT scaffolds, with a dense cell layer located at the periphery. Blue highlights DAPI labeled nuclei, as well as autofluorescence of the DAT scaffold. Higher magnification image of boxed region is shown on right. Scale bar is 1 mm. (b) At endpoint, the implants were dissected and processed for flow cytometric analysis. Quantification of the DsRED^+^ cells in the seeded implants highlights the decline in this population over time (n= 5-6 mice per timepoint with at least 2 mice of each sex per timepoint).

### 3.2 ASC phenotype is minimally impacted by implantation

To better understand the rapid clearance of the donor ASCs, a mass spectrometry-based proteomics approach was employed to characterize the phenotype of the DsRED^+^ ASCs isolated from the DAT implants in the *in vivo* study at 3- and 7-days post-implantation in comparison to ASCs grown on TCPS or the DAT scaffolds prior to implantation. Following the selection criteria, a total of 2585 unique proteins were identified. Marked differences existed between the cells cultured on TCPS and those that were extracted from any of the DAT-seeded bioscaffold groups. As expected, several contractile proteins including calponin 1, transgelin, alpha actinin, gamma 1 actin, alpha cardiac muscle actin, and myosin light chain 9, were expressed at greater levels in the cells grown on TCPS. Interestingly, while the TCPS samples expressed higher amounts of type V collagen, types IV and VI collagens were more abundant in the DAT-seeded groups. Several Aldehyde Dehydrogenase (ALDH) proteins, including 1l1, 6a1, 9a1, 1a2, 4a1, and 3a2, were upregulated in the DAT-seeded group. Moreover, cellular retinoic acid binding protein 1 (Crabp1), a marker that has been used to identify a more regenerative fibroblast population within skin^37^ was expressed over 3-fold greater in the combined DAT-seeded ASCs compared to the TCPS controls. Overall, 370 proteins were expressed at significantly greater amounts in DAT-seeded ASCs, whereas 115 were relatively over-expressed on TCPS (the entire dataset is available in the included Supplemental Data File). Within the DAT-seeded groups, differences were only noted between the 3-day post-implantation group and the non-implanted controls where 6 proteins were relatively upregulated in the 3-day samples, and 11 proteins were relatively upregulated in the non-implanted DAT-seeded samples. Notably, none of these differentially expressed proteins are classically associated with apoptosis, indicating that it is unlikely that this is a major mechanism driving the clearance of the ASC population. These data can be further visualized with the hierarchical clustering and principal component analyses (PCA), highlighting the similarities between all DAT-seeded samples, and the contrast with ASCs cultured on TCPS (Figure 2). Although no differentially expressed proteins were detected between the day 3 and day 7 implants, there is a notable difference between these samples along the component 2 axis of the PCA, suggesting that holistically, there is a shift in this population over time. Additionally, one of the non-implanted DAT-seeded ASC samples was more like the TCPS control samples. This may be due to natural heterogeneity within the DAT scaffolds, as more fibrous regions could potentially promote a more fibrotic phenotype within the ASCs.

**Figure 2:**
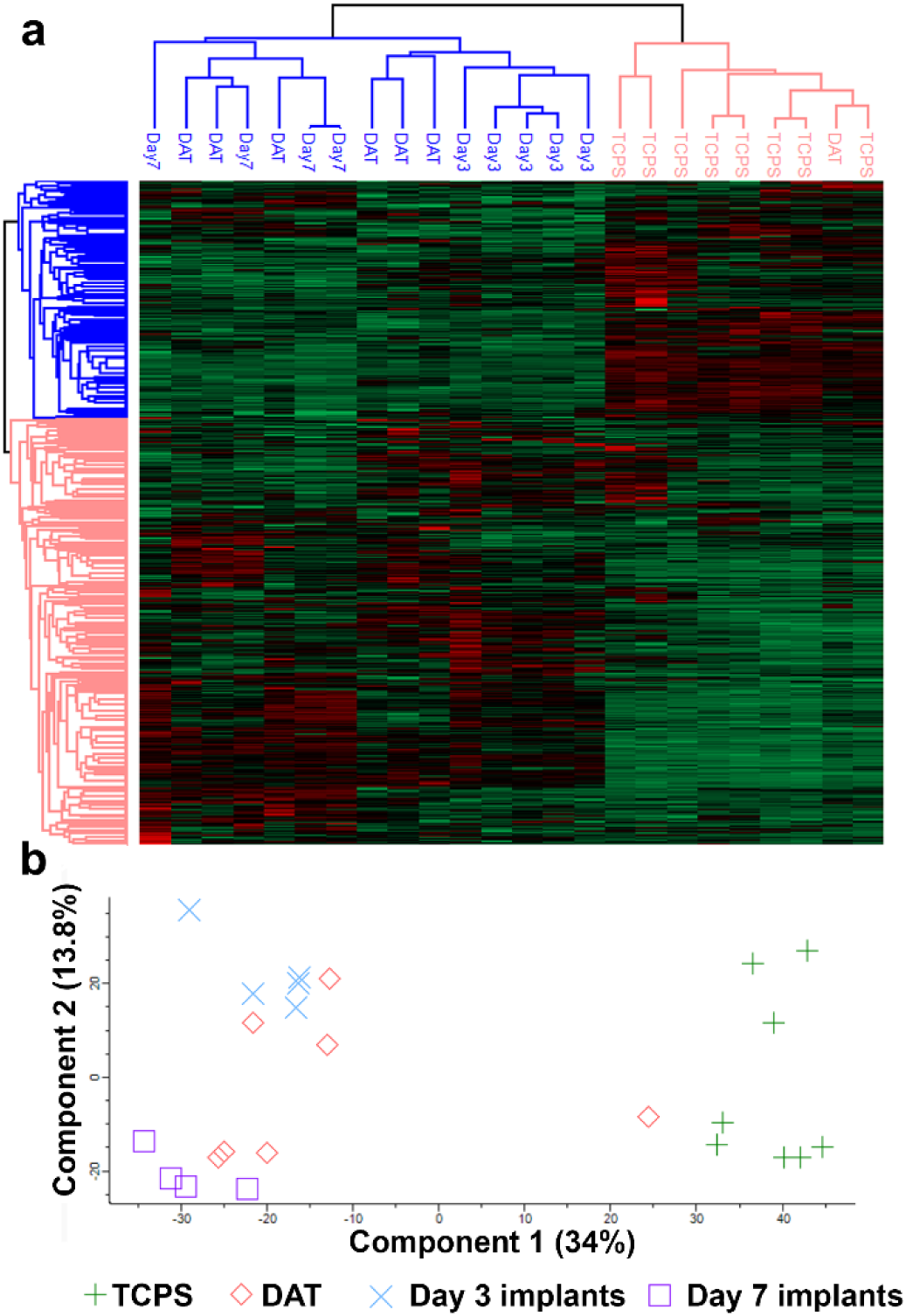
ASC phenotype is primarily defined by substrate, with similar expression patterns observed in the ASCs on the DAT scaffolds both pre- and post-implantation. ASC-seeded scaffolds were isolated and digested at endpoint and ASCs were further purified via FACS. Sorted ASCs were then lysed, and the proteome was assessed with mass spectrometry. Samples up to 7 days post-implantation were included for further analysis. (a) Hierarchical clustering and (b) principal component analysis identified two major populations of ASCs defined by their adherence to culture plastic (TCPS) or the DAT biomaterials (n = 4-8 samples per group; female donor cells and recipient mice were used).

### 3.3 Donor ASCs modulate the macrophage population at the implant site

With limited change in ASC phenotype post-implantation, it was expected that their well-characterized immunomodulatory phenotype would be preserved following delivery. To investigate the effects that ASCs had on macrophage polarization, implants were harvested and digested for analysis via flow cytometry, comparing the seeded and unseeded DAT scaffolds (Figure 3a). Macrophages were characterized based on their expression of CD11b and F4/80, and were further probed for their expression of CD163, CD68, Arginase I, and iNOS (figure 3b). Macrophages in the seeded group had a significantly greater percent positivity for Arginase I and CD68 (Figure 3b; main effect of seeding p<0.05; n =5-6 mice per timepoint), with overall similar levels of iNOS and CD163 (Figure 3c). Importantly, the noted differences were greatest within the first week post-implantation when the ASCs were most abundant.

**Figure 3:**
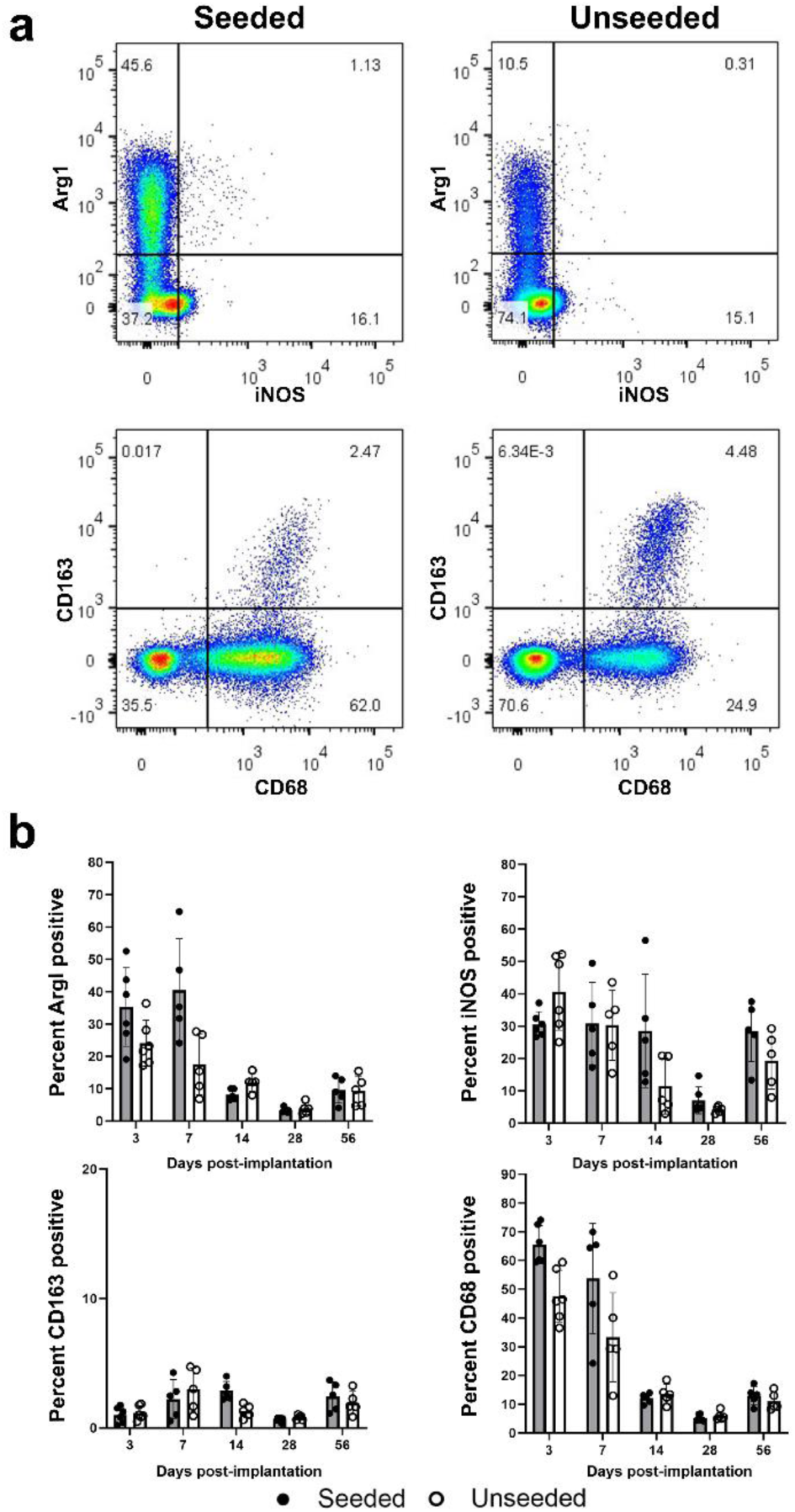
ASC seeding of DAT increases the percentage of host macrophages expressing Arginase I and CD68 at the implant site. (a) Scaffolds were removed at endpoint and digested for cell isolation and analysis via flow cytometry. Representative flow plots are shown from 7-day implants. (b) Quantification of macrophage markers showed significant main effects for seeding on the expression of Arginase I and CD68 within the macrophage population (linear mixed effects model; p < 0.05). Seeding did not affect macrophage positivity for CD163 or iNOS. n= 5-6 mice per timepoint with at least 2 mice of each sex per timepoint.

### 3.4 Host macrophages phagocytose donor ASCs

With minimal changes detected in the ASC phenotype within the DAT implants over time, we hypothesized that ASC clearance was not primarily mediated by apoptosis. Notably, their depletion from the implant site was correlated with increased presence of CD45^+^ bone marrow-derived populations (Figure 4a). Due to the early appearance of these cells at the implant site, it was hypothesized that many of these were likely macrophages. To further explore their identity, tissues were probed with an antibody for IBA1, a macrophage specific protein involved in phagocytosis^38^. Confocal microscopy at the wound edge confirmed that many of these CD45^+^ cells were indeed macrophages, and interestingly, some of these appeared to have phagocytosed DsRED protein. Further analysis via flow cytometry supported this finding. Specifically, a small population of DsRED^+^CD11b^+^F4/80^+^ macrophages was observed. Recent data have identified phagocytosis of mesenchymal stromal cell (MSC) populations by macrophages as a driver of macrophage polarization^39–41^. To explore whether this might be relevant in this model, macrophage expression of Arginase I, iNOS, CD163, and CD68 was compared between DsRED^+^ macrophages and their DsRED^−^ counterparts within the seeded implant group (Figure 4c). Interestingly, Arginase I, CD163, and CD68 were all expressed in a greater proportion of the DsRED^+^ macrophage population, whereas iNOS was expressed in a greater proportion of the DsRED^−^ macrophage population (main effect DsRED^+^ vs. DsRED^−^ macrophages: p<0.05 for all markers; Figure 4d).

**Figure 4:**
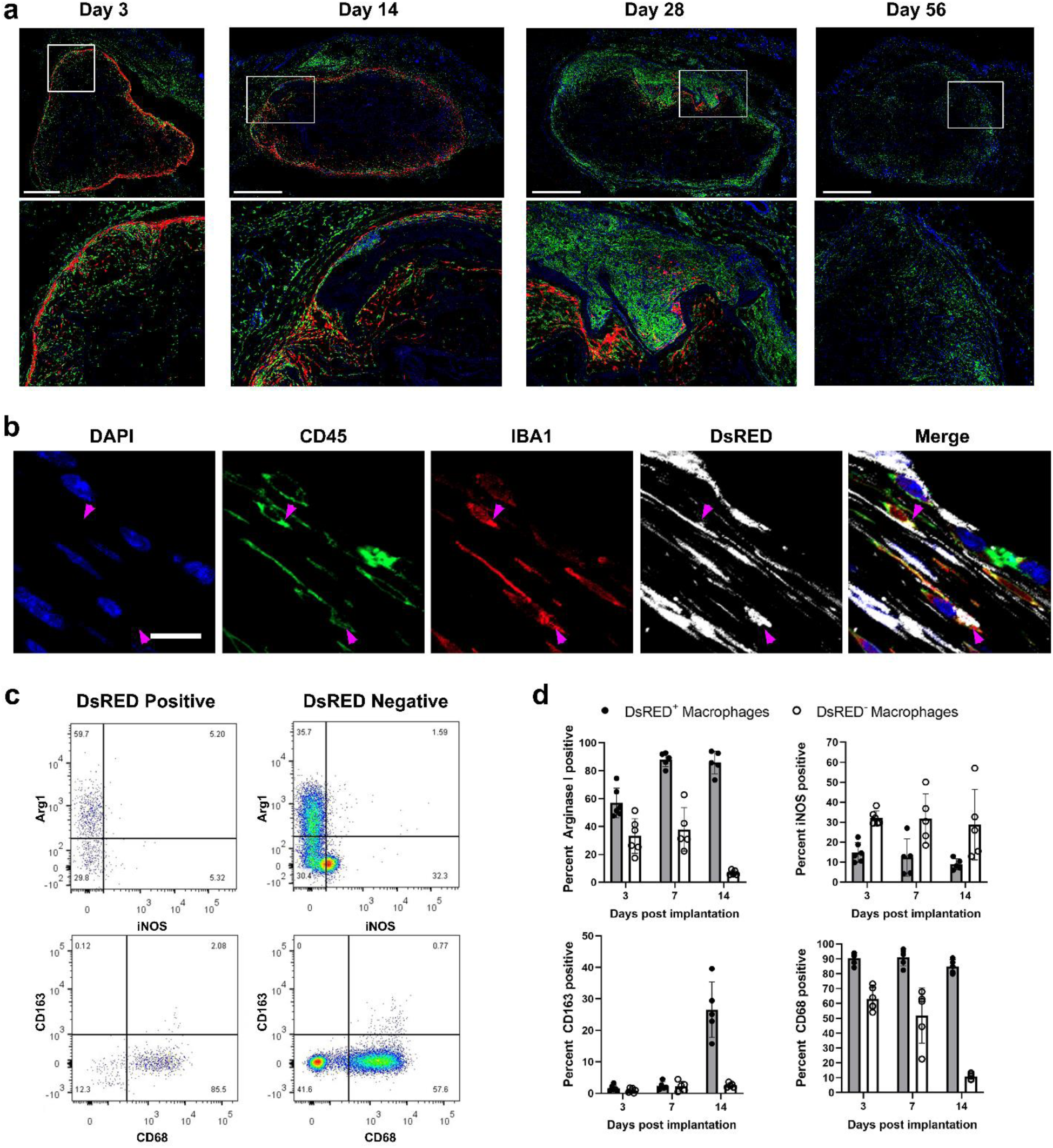
ASCs are cleared from the implant site by macrophages that display a unique phenotype compared to the bulk macrophage population. (a) IF assessment of CD45 (green) and DsRED (red) in ASC-seeded implants shows that the loss of ASCs over time correlates with CD45^+^ cell infiltration. DAPI labeling in blue. Scale bar is 1 mm; marked insets shown below. (b) Confocal microscopy at the outer edge of ASC-seeded implants at day 7 identified macrophages that have phagocytosed DsRED protein (marked by the magenta arrows; scale bar is 10 µm). (c,d) In the flow cytometric analysis of these implants, a subset of DsRED^+^ macrophages were observed to have an altered expression profile compared to the DsRED^−^ population, showing enhanced expression of CD68, as well as the pro-regenerative markers Arg-1 and CD163, and reduced iNOS expression. Representative flow plots are shown from day 3 samples. Significant main effect of scaffold seeding for all markers (linear mixed effects model; p < 0.05). n= 5-6 mice per timepoint with at least 2 mice of each sex per timepoint.

### 3.5 ASC-seeding does not affect the perfused vascular network detected at the implant site at 4 and 8 weeks

To examine whether seeding with ASCs modified the angiogenic response, a micro-CT-based angiography method was employed to visualize the vascular network in the DAT implants in mice at 28- and 56-days post-implantation. Qualitatively, the scans showed no obvious differences in vessel structure or density (Figure 5a). This was further supported by quantitatively assessing voxel scalar values taken for the entire scaffold volume and the 200 µm surrounding the scaffold (Figure 5b). From the µCT analysis, the scaffold size measured in the 56-day cohort was significantly smaller than those measured at 28-days post-implantation, suggesting that the scaffolds were being remodeled over time (Supplemental Figure 1).

**Figure 5:**
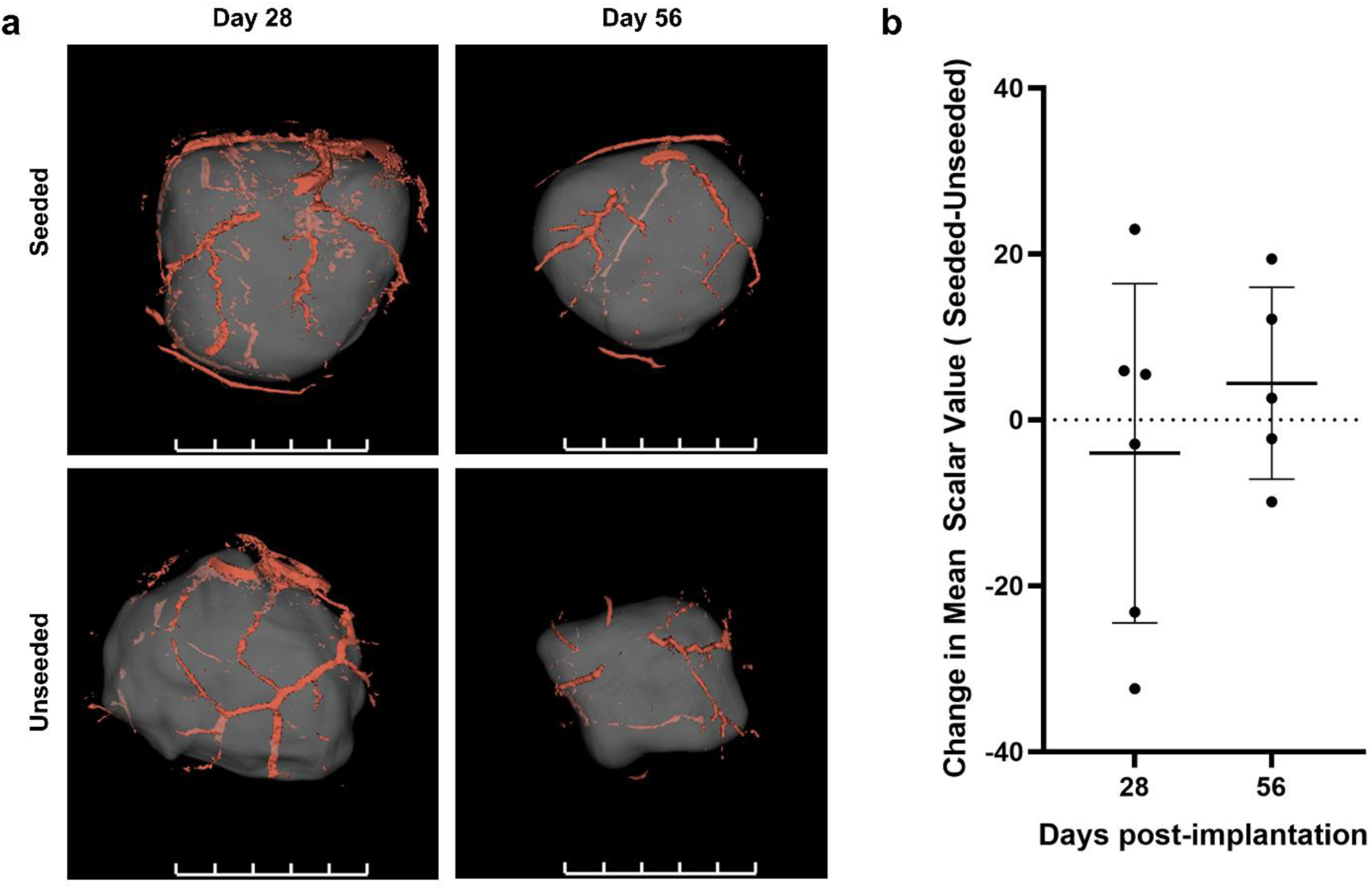
Scaffold seeding did not affect the density of the perfused vascular network at the implant site. (a) µCT-based angiography was used to investigate the perfused vascular network at the implant site. 3D models were derived from these data to highlight the vasculature within 0.5 mm of DAT implant surface, showing qualitatively similar vascular networks. (b) The amount of contrast agent within the scaffold volume was quantitatively assessed by averaging the scalar value of each voxel within 200 µm of the scaffold surface. Differences in this value between seeded and unseeded scaffolds were centered around 0, suggesting that similar amounts of contrast agent were present between groups (p > 0.05; n = 5-6 mice per timepoint with a minimum of 2 of each sex per timepoint).

### 3.6 Both ASC-seeded and unseeded DAT implants were well integrated with the host tissues at 8 weeks

Finally, IF staining of the implants at the terminal endpoint of 8 weeks suggested that both ASC-seeded and unseeded DAT implants integrated well with the host tissues. Specifically, perilipin staining showed that the adipocytes in both groups remained directly adjacent to the implanted scaffolds (Figure 6a). Although endogenous fibroblasts, identified by PDGFRα expression, were located at the edge of the implants, there was no evidence of a fibrous capsule forming around the scaffolds, suggesting that both groups were well tolerated. CD31^+^ vascular structures were observed within the implants in both groups, although these were typically small vascular structures without notable αSMA^+^ mural cell support (Figure 6b). However, the tissue surrounding the implants contained αSMA^+^ mural cell-supported vessels, consistent with healthy neighboring host tissue. Overall, these findings provide evidence of long-term and well-tolerated integration of the scaffolds from both groups within the host tissue.

**Figure 6:**
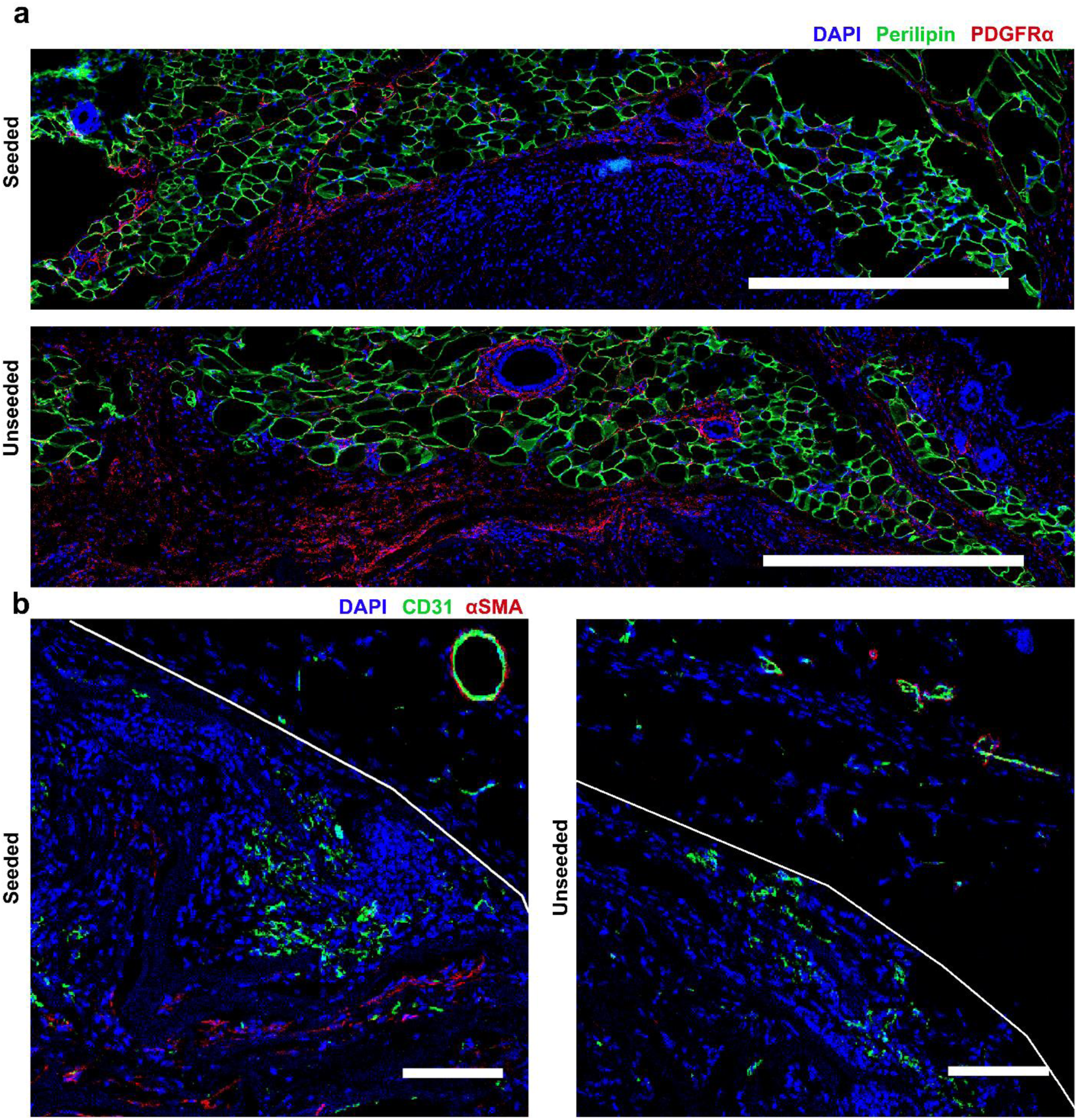
Long-term regenerative outcomes are similar between seeded and unseeded scaffolds. At 8-weeks post implantation, (a) IF assessment of perilipin showed adipocytes from the neighboring fat pad surrounding both scaffold groups (scale bar is 500 µm). (b) Similarly, staining for CD31 and α-smooth muscle actin showed that most of the vascular structures within the implant were small and not supported by mural cells (scale bar is 100 µm; white lines indicate the scaffold edge)

## 3. Discussion

Adipose tissue is an important endocrine organ with critical functions throughout life and is essential for maintaining the normal contours of the human body. Despite advances in plastic and reconstructive surgeries, the regeneration of stable adipose tissue following subcutaneous tissue loss remains limited. Thus, there remains a need to develop strategies that can promote adipose tissue regeneration to enable long-term large volume augmentation. Previously, we have identified DAT as a promising adipo-conductive material, which supports the development of new host-derived fat when implanted subcutaneously in animal models^12,13,23,42,43^. ASCs cultured *in vitro* on DAT display enhanced adipogenic, angiogenic, and immunomodulatory signaling^15,21^ and when combined, ASC-seeded DAT implants have shown enhanced therapeutic efficacy in preclinical models compared to DAT alone^12,13,42,43^. With the long-term goal of enhancing the pro-regenerative capacity of this platform, this study aimed to better understand the mechanisms through which ASCs mediate *in vivo* adipose tissue regeneration using syngeneic donor cells in an immune competent model.

We have previously observed allogeneic rat ASCs from male donors persisting within DAT scaffolds implanted subcutaneously within the dorsal skin of immune competent female Wistar rats for up to 8-weeks post-implantation by fluorescence *in situ* hybridization analysis of the y-chromosome^13^. Similarly, using a syngeneic mouse model to investigate periodontal regeneration, Lemaitre *et al.* (2017) implanted mouse ASCs within a collagen gel and observed histologically that donor cells persisted up to 6-weeks post-implantation, but were not present at 12 weeks^44^. In the present study, the syngeneic donor ASCs were rarely observed within tissue sections collected at 8-weeks post-implantation in the inguinal implant model. Both IF and flow cytometric assessment supported that the donor cells were mostly cleared prior to 4-weeks post-implantation. Notably, the imaging results highlighted that the DsRED^+^ cells were depleted from the outer regions of the implants at earlier timepoints, followed by the cells that resided within the interior of the implant, which correlated with a progression in host CD45^+^ cell infiltration.

To our knowledge, quantitative flow cytometric data on the abundance of delivered ASCs in immune competent animals has not previously been reported. Using a bioluminescence reporter to track human ASCs in dorsal subcutaneous DAT-based implants within immunocompromised nude mice, Morissette-Martin *et al.* (2021) showed a similar trajectory, with ASCs decreasing in abundance over the first 2-weeks post-implantation. However, in this model, the bioluminescence signal plateaued thereafter, up to 4-weeks post-implantation^29^. Similarly, Suga *et al.* (2014) observed a rapid decline in mouse ASC abundance via bioluminescence imaging over two weeks following direct injection within PBS into the inguinal fat pad of immune competent mice^28^. Additionally, Todeschi *et al.* (2015) observed the clearance of umbilical-derived, but not bone marrow-derived MSCs, within 30 days following dorsal subcutaneous implantation on a ceramic scaffold using a bioluminescence reporter system^31^. Using PCR to track human ASCs injected in suspension or as spheroids into the inguinal region of either immune competent or athymic nude mice, Agrawal *et al.* (2014) were unable to detect donor cells by 3-weeks within the immune competent model, compared to about 5% of the initial signal detected within the immunocompromised mice by this timepoint^30^. Notably, within both systems, the majority of cells were lost within the first 10 days post-implantation. Thus, even within these disparate subcutaneous implant models, rapid clearance of ASCs is common. Importantly, recognizing that the cell dose delivered can be an important mediator of the *in vivo* regenerative outcomes^45,46^, this rapid clearance is likely limiting the potential therapeutic benefit of the ASCs. In the present study, short-term modulation of macrophage phenotype was noted in the ASC-seeded groups, but it was evident that the ASCs had limited long-term effects on the regenerative outcomes.

The mechanisms through which ASCs are cleared from tissues are not completely understood. As mentioned above, in the current study, the loss of ASCs correlated temporally and spatially with the influx of CD45^+^ cells into the DAT implants. Furthermore, DsRED^+^ macrophages were observed via IF staining and flow cytometry starting at the earliest timepoint of 3 days. Still, it is uncertain whether the ASCs undergo apoptosis in response to environmental conditions and are subsequently cleared by macrophages and/or if they are targeted for clearance by host immune cells. Proteomic profiling of the ASCs showed little change in their expression patterns up to 7-days post-implantation, compared to the ASCs extracted from the DAT implants immediately prior to implantation, with no evidence to indicate a shift towards apoptosis. However, it is important to acknowledge that the proteomics profile includes the most abundant proteins, which limits the resolution at which we can understand their phenotype. Interestingly, Suga *et al* (2014) showed greater persistence of implanted ASCs via bioluminescence tracking within ischemic inguinal fat pads relative to control fat pads^28^, which could be indicative of circulating immune cells playing a critical role in ASC clearance.

It was unexpected that the ASC phenotype was similar following implantation relative to ASCs isolated from seeded scaffolds prior to implantation. Notably, all groups were considerably different compared to those cultured on plastic, which is likely related at least in part to differences in mechanical stiffness, a phenomenon that has been well described by others^47–50^. The similarity between the ASCs cultured on the DAT scaffolds pre- and post-implantation is an important finding as it supports the use of these biomaterials as soft substrates for characterization of ASCs *in vitro* with relevance to their behavior *in vivo*. Nevertheless, it is possible that there are more subtle differences between these samples that we were not able to detect with this method. More sensitive approaches such as single cell RNA-sequencing (scRNA-seq) may be able to provide a deeper understanding of ASC behavior post-implantation. For example, when observing the PCA from the proteomics data, it is evident that global expression patterns of the ASCs may be changing over time following implantation. Performing a pseudotime analysis of scRNA-seq data following induced muscle injury, Scott *et al.* (2019) showed the extent to which several endogenous mesenchymal populations, including ASC-like fibro-adipogenic progenitors (FAPs), change over time alongside the inflammatory and healing processes^51^. Interestingly, this endogenous population undergoes activation following injury and transitions back into its homeostatic state following resolution. It would be expected that, following implantation, the resultant innate immune response would be similar and thus the ASCs would be susceptible to similar changes over time. Further understanding of how the phenotype of ASCs changes following implantation will give critical information on their function over time and may also give insight into the mechanisms through which they are cleared.

Notably, others have reported that apoptosis of MSCs and their subsequent phagocytosis are key mechanisms through which they can mediate macrophage polarization^39–41^. In this study, phagocytic DsRED^+^ macrophages showed greater positivity for Arginase I, CD163, and CD68 compared to DsRED^−^ macrophages within the same tissue, while simultaneously showing decreased positivity for iNOS. The confocal microscopy images reveal that these phagocytic macrophages are in close proximity with the transplanted ASCs, supporting that these expression changes may be mediated by crosstalk between the two populations. However, we cannot rule out the possibility that the DsRED^+^ macrophages simply represent a more phagocytic subpopulation of macrophages with an alternative phenotype.

It is worth noting that a limitation of tracking cells *in vivo* using exogenous proteins, such as fluorescent or bioluminescent proteins, is that the proteins themselves can stimulate an immune response affecting the persistence of the labelled cells^52–55^. However, without ASC-specific markers for identification, use of these strategies is a cost-effective approach to track cells following implantation that has been widely adopted. Nevertheless, in the context of regeneration, there is little known about how these reporter proteins affect long-term cell survival or regenerative outcomes, which is a critical area of investigation for future studies.

Investigation of long-term regenerative outcomes suggested a high level of similarity between ASC-seeded and unseeded scaffolds up to 8-weeks post-implantation. Despite early changes in macrophage phenotype, once the ASCs were largely depleted from the implants after 2 weeks, macrophage marker expression normalized between the treatment groups. Similarly, implant vascularization, scaffold remodeling, and overall integration with the host tissues were similar, independent of seeding. It is evident from micro-CT analysis that the scaffolds are shrinking in size over time, but whether this is due to remodeling into *de novo* adipose tissue, or simply the result of degradation of the scaffold is difficult to conclude with the methods used in this study. Despite delivering large numbers of cells, these results are in contrast with those of previously published work, potentially due to model selection and the methods used to quantify these parameters. For example, several studies have used histological quantification of CD31^+^ structures to show that MSC-seeded biomaterials are more angiogenic than unseeded controls^12,13,29,31,56^. Albeit with less resolution, the angiography method employed in the present study showed no changes qualitatively or quantitatively to the perfused vascular network between the seeded and unseeded groups. It has been well characterized that vascularization following inflammation leads to the rapid influx of poorly perfused vessels^34^, a process that has been hypothesized to promote fibrotic outcomes^57^. Thus, it is not surprising to see an increase in vessel density following implantation. More importantly, it is unclear whether the influx of poorly perfused vessels is predictive of improved outcomes. In previous work, we explored both histological vessel density, and vascular perfusion using micro-CT-based angiography following implantation of human ASCs delivered on a DAT scaffold into immunocompromised nude mice^29^. Notably, in that study, increased vessel density measured histologically did not correlate with increased vascular perfusion. Further work correlating vessel density with vascular perfusion could help to better understand how these parameters affect regenerative outcomes.

Overall, the results highlight an inherent challenge with cell-based approaches, which is that infiltrating host immune cell populations that are required for implant remodeling and tissue regeneration will also clear the therapeutic cell population. Increasing the initial therapeutic cell dose may not be sufficient and may in fact expedite host immune cell infiltration and subsequent ASC clearance, although increasing the ASC density in the interior of the scaffolds may be advantageous to have more sustained effects. A potential strategy that would be interesting to explore would be to augment the donor ASC population through localized cell injection into the implant region at 1-2 weeks post-implantation, to see if this could enhance the long-term regenerative response.

## 4. Conclusions

Advanced biological characterization methods were used to assess the effects of syngeneic ASC seeding on regenerative outcomes in DAT bioscaffolds implanted into the inguinal region of immunocompetent C57BL/6 mice. Interestingly, mass spectrometry-based proteomics revealed that the phenotype of ASCs cultured or delivered on the DAT biomaterials was markedly different compared to ASCs cultured on the TCPS, however, *in vivo* implantation did not substantially alter the ASC phenotype compared to the pre-implantation scaffold controls. These findings support that the *in vitro* characterization of ASC-seeded DAT biomaterials is relevant for understanding their behavior *in vivo*. While the ASC-seeded DAT scaffolds modulated infiltrating host macrophage polarization at early timepoints, there were no notable differences in macrophage phenotype, vascular perfusion, implant remodeling, or overall integration over the longer-term compared to unseeded DAT scaffolds. The limited effects of ASC delivery in this model are likely attributed to the rapid depletion of the donor cells, suggesting that strategies to promote longer-term retention or replenishment of the therapeutic ASC population should be the focus of further research.

## Supporting information

Supplemental data 2 - Proteomics

## Acknowledgements

The authors would like to thank Dr. Damir Matic and the teams of Dr. Aaron Grant and Dr. Damir Matic for providing the adipose samples used in this study. Further, we thank Dr. Kristin Chadwick and the London Regional Flow Cytometry Facility for guidance and assistance with flow cytometry and cell sorting experiments. This work was funded by the Canadian Institutes of Health Research (CIHR; FRN# 119394 and FRN# 179858), with salary support provided by the CONNECT! NSERC Collaborative Research and Training Experience Program (CREATE) in Soft Connective Tissue Regeneration/Therapy.

