## Supplemental data 1 - Figures and Tables for "Syngeneic adipose-derived stromal cells modulate the immune response but have limited persistence within decellularized adipose tissue implants in C57BL/6 mice"

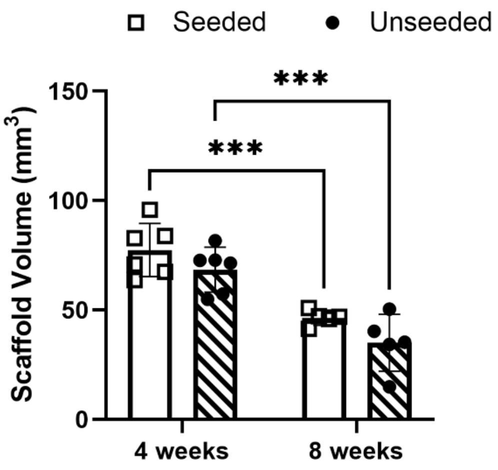


**Figure S1:** Scaffold volume decreased over time. Micro-CT scans were performed at 4- and 8-week endpoints, and the scaffolds were manually segmented out in 3D slicer software. No differences were detected between the scaffold groups, but were noted across timepoints within groups ($p < 0.001$). n= 5-6 mice per timepoint.

**Supplemental Table 1:** Primary antibodies used for immunofluorescence labelling of mouse tissues. Antibodies within the same staining pair were assessed simultaneously.

| Target | Species | Staining pair | Supplier (catalog number) | Dilution |
|---|---|---|---|---|
| αSMA | Rabbit | 1 | Abcam (Ab5694) | 1:100 |
| CD31 | Goat | 1 | BioTechne (AF3628) | 1:50 |
| CD45 | Goat | 2 | BioTechne (AF114) | 1:100 |
| IBA1 | Rabbit | 2 | Abcam (Ab178846) | 1:100 |
| PDGFRα | Goat | 3 | BioTechne (AF1062) | 1:100 |
| Perilipin | Rabbit | 3 | New England Biolabs (9349S) | 1:200 |

**Supplemental Table 2:** Secondary antibodies used for immunofluorescence labelling of mouse tissues.

| Target species | Conjugated fluorophore | Supplier (catalog number) | Dilution |
|---|---|---|---|
| Goat | Alexa Fluor 488 | Thermo Fisher Scientific (A-11055) | 1:500 |
| Goat | Alexa Fluor 647 | Thermo Fisher Scientific (A-21447) | 1:500 |
| Rabbit | Alexa Fluor 488 | Thermo Fisher Scientific (A-21206) | 1:500 |
| Rabbit | Alexa Fluor 647 | Thermo Fisher Scientific (A-31573) | 1:500 |

**Supplemental table 3:** Antibodies used for fluorescence activated cell sorting to isolate donor ASCs.

| Target (or label) | Fluorophore | Supplier (catalog number) | Dilution | Laser and filter set |
|---|---|---|---|---|
| CD11b | PE-Cy5 | BioLegend (101210) | 1:160 | Yellow-Green 670/14 |
| SYTOX Blue | | Thermo Fisher Scientific (S34857) | 1 µM | Violet 450/40 |
| DsRED | | | | Yellow-Green 582/15 |

**Supplemental table 4:** Data acquisition parameters for Q Exactive Plus mass spectrometer.

| **Parameters** | **QE Plus Orbitrap** |
|---|---|
| Mass Range (m/z) | 300-1400 |
| Isolation Window (m/z) | 1.3 |
| MS Resolution | 70K @ 200 m/z |
| MSMS Resolution | 17.5K |
| MS Injection Time (ms) | 250 |
| AGC Target (MS) | 1E6 |
| AGC (Target MSn) | 2E5 |
| Preview Scan | n/a |
| Threshold (counts) | 50K |
| Underfill Ratio | 3% |
| Data Dependent Acquisition | Top 10 |
| Dynamic Exclusion (s) | 30 |
| Exclusion Mass Width (m/z) | n/a |
| Exclude Isotopes/Monoisotopic Precursor Selection | Enabled |
| Fragmentation Type | HCD |
| Normalized Collison Energy | 27 |
| Lock Mass (445.120025m/z) | Best |
| Charge State Rejection | Unassigned and +1 |
| Default Charge State | +2 |

**Supplemental table 5:** Antibodies used for flow cytometry analysis of donor ASCs and host macrophages.

| **Target (or label)** | **Fluorophore** | **Supplier (catalog number)** | **Dilution** | **Laser and filter set** |
|---|---|---|---|---|
| CD68 | Brilliant Violet 605 | BioLegend (137021) | 1:50 | Violet 610/20 |
| F4/80 | Alexa Fluor 488 | Thermo Fisher Scientific (53-4801-82) | 1:80 | Blue 530/30 |
| CD11b | PE-Cy5 | BioLegend (101210) | 1:160 | Yellow-green 670/14 |
| Arginase I | PE-Cy7 | Thermo Fisher Scientific (25-3697-82) | 1:200 | Yellow-green 780/60 |
| CD163 | APC | Thermo Fisher Scientific (17-1631-82) | 1:200 | Red 670/30 |
| iNOS | APC-eFluor780 | Thermo Fisher Scientific (47-5920-82) | 1:150 | Red 780/60 |
| LIVE/DEAD™ Fixable aqua | | | | Violet 525/50 |
| DsRED | | | | Yellow-green 582/15 |
